# Complex dynamics arising from the interplay between prey harvest saturation and predator-prey dynamics

**DOI:** 10.1101/2020.09.22.309203

**Authors:** Michel Iskin da S. Costa, Lucas dos Anjos

**Affiliations:** Laboratório Nacional de Computação Científica, Av. Getúlio Vargas, 333 – Quitandinha, Petrópolis, RJ 25651–070 Brazil

**Author notes:** Corresponding author. E–mail addresses (M.I.S. Costa), (L. dos Anjos).

**Keywords:** Prey harvest saturation, predator-prey, complex dynamics, management policies

## Abstract

A study of the influence of prey harvest saturation on the dynamics of a predator-prey system is undertaken. It is shown that the augmentation of the constant intensity harvest part of this functional response can significantly change prey-predator dynamics by means of slight variations of the constant harvest intensity, rendering thus species management difficult due to, for instance, multiple attractors. Given that some management policies rely on the harvest saturation structure studied in this work, these results may have significant implications regarding renewable resource management.

## 1 Introduction

Harvesting or any kind of species removal has been a subject of theoretical studies in population as well as in community dynamics. For instance, species overfishing has triggered many severe and unexpected behaviors such as communities that are more prone to cycles of booms and busts and as a consequence, these sudden and unexpected shifts can render the fishery system less manageable (Travis et al., 2014). By means of numerical simulations, in this work we show that unexpected shifts leading to complex dynamics may arise in a relatively simple predator-prey model where prey is the only harvested species. Given that some management policies are based on the harvest function studied in this work, the dynamical outcomes found here may have significant implications regarding renewable resource management.

## 2 A predator-prey model with prey harvest saturation

In terms of renewable resource management, harvest saturation consists of a fixed limit of harvest capacity. More precisely, suppose that a harvesting procedure is capable of capturing a maximal quantity of biomass (or individuals) *C* per unit time. Below this maximum capacity the capture rate is assumed to be proportional to the density of the exploited stock up to the maximum capacity level *C*. Beyond this level, capture rate is assumed to be constant and equal to *C*. Denoting *h* as the harvest rate of a population *x*, harvest saturation can be formulated as (Clark, 1976):

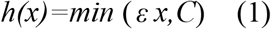

where *ε* is the harvest effort. For 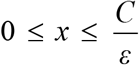 it is said that the capture is undertaken in a constant harvest effort fashion while for 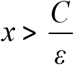, under a constant harvest rate fashion.

The trophic scheme to be analyzed in this work is displayed in figure 1.

**Figure 1.**
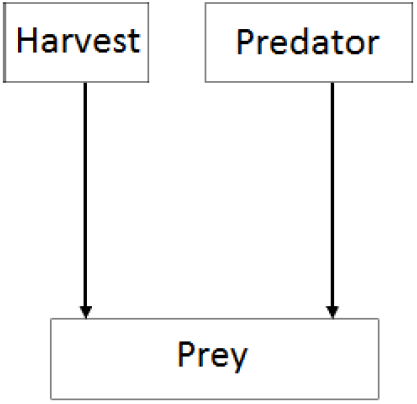
A diagram of a predator-prey system with prey harvest. Arrows denote consumption/extraction.

Figure 1 depicts a trophic scheme where prey is simultaneously harvested and preyed upon by a predator. This food web diagram was proposed by Yodzis (2001) in order to assess the influence of predator culling (removal) on prey fishing yield (the so-called ‘surplus yield’ hypothesis).

A continuous time dynamical model describing the behavior of the species involved in the trophic scheme of figure 1 can be given by:

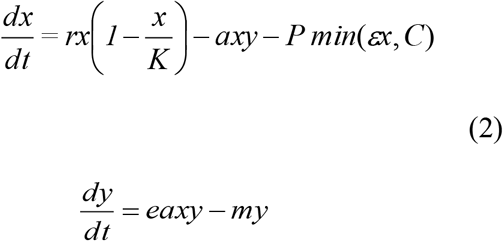

*x* and *y* are the prey and specialist predator populations, respectively; *r* is the prey specific growth rate and *K* is its carrying capacity; *a* is a predation attack coefficient of a functional response type 1, and *e* is the predator food–to–offspring conversion coefficient; *m* is the density independent per capita mortality rate of the predator; *h(x)=min*(*εx, C*) is defined in (1) where *ε* is the constant harvest effort on prey and *C* is the maximal harvest capacity. In terms of prey harvest, the constant *P* could be ascribed, for instance, to a fleet consisting of a constant number of fishery vessels in the dynamical equation of the prey *x* (without loss of generality we consider *P*=1).

System (2) consists of two structures, namely:

**Linear proportional harvest**

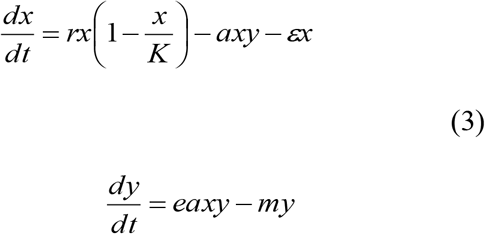
**Fixed harvest rate (constant harvest rate)**

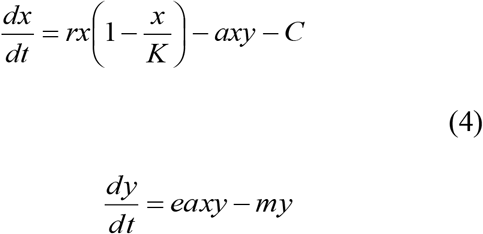

Structures (3) and (4) correspond to two prey-predator models which are separated in the plane *x vs. y* by the line 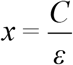 (note that the constant harvest effort structure (3) is actually a Volterra model (Turchin, 2003) with prey being consumed by a dynamical predator (*y*) with functional response type 1 given by the term *ax*; the additional predation term *εx* can be incorporated in the logistic term as 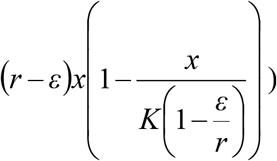. Accordingly, the phase plane *x vs. y* is split into two regions – one for constant harvest effort (left to the line 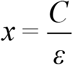 – region *PH*) and other for fixed harvest rate (right to the line 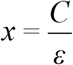 – region *FH*) (see figure 2; the terms *PH* and *FH* will be used throughout the text). Since each structure possesses its own nontrivial equilibrium points, the position of the line 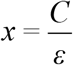 (sometimes denoted switching line) will determine whether the points lie in their respective region or not. In the first case they are called real equilibrium points, while in the second they are named virtual equilibrium points (Costa et al., 2000).

**Figure 2.**
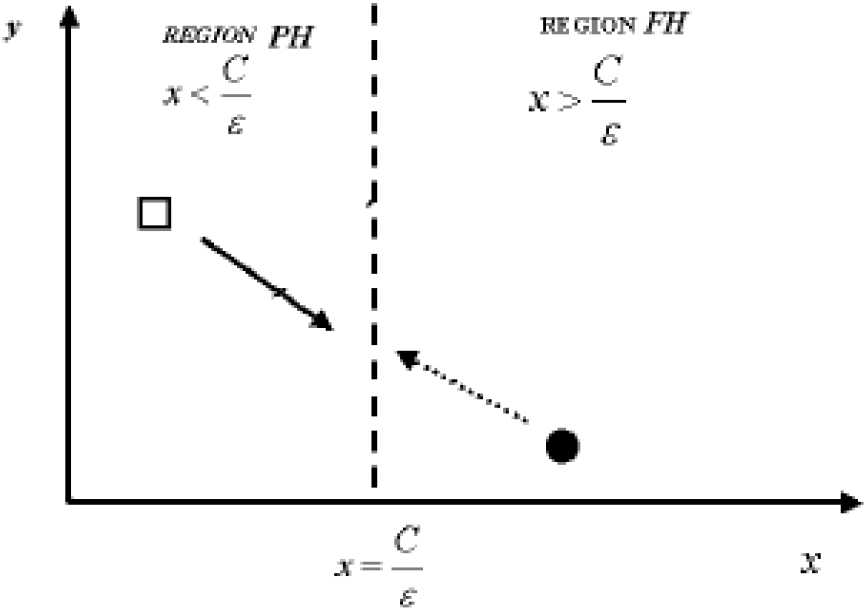
A schematic figure of the two structures in the plane *y vs. x*.

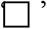 – equilibrium point of region 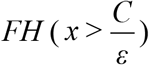. 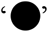 – equilibrium point of region 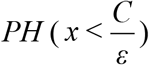. These points are not located in their respective regions. Hence they are virtual equilibrium points. The solid arrow indicates the vector field directed to the constant harvest rate equilibrium point, while the dashed arrow indicates the vector field directed to the constant harvest effort (or proportional harvest) equilibrium point.

In this work, we focus on the effects of prey harvest saturation (*C*) on the dynamics of system (2). Hence, the following analysis will consist of drawing phase planes of this system for increasing values of *C*. In terms of harvest of renewable resources, this mathematical setup could well describe the expansion of harvest capacity in trawl fishery. Keeping the fleet size constant (number of fishery vessels), one could replace the same number of boats with new ones with a larger capacity (processing a higher number/biomass of fish per time unit, i.e., a higher *C*). This procedure could be undertaken maintaining the same mesh size in all vessels as before (i.e., keeping the same value of the fishing effort *ε*).

According to model (2), augmentation of *C* will push downwards the *x* isocline *C* (*dx/dt*=0 – a parabola in the *FH* region) while the vertical line 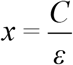 will slide horizontally rightwards along the prey *x*-axis. Next we show phase planes of model (2) for increasing values of *C*.

Figure 3 displays a phase plane of model (2) for *C*=0.1.

**Figure 3.**
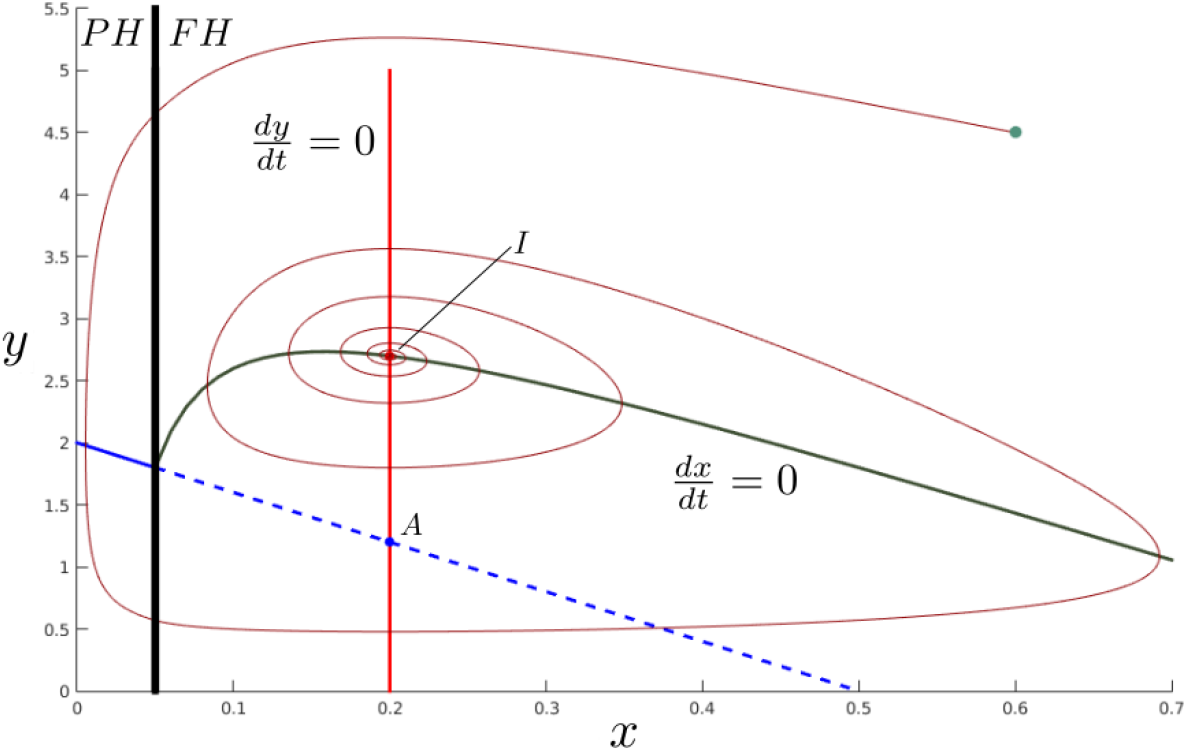
One phase plane of model (2) for *C* = 0.1. The trajectories converge to the sole real and stable equilibrium point *I* of the fixed harvest (*FH*) region. Point *A* of the *PH* region is virtual. 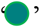 – initial conditions. Parameter values: *r*=1; *K*=1; *a*=1; *e*=2.5; *m*=0.5; *ε* =0.05; *C*=0.1;

The simulated trajectory converges to the sole real and stable equilibrium point *I* of the fixed harvest (*FH*) region. Note that the point *A* of the *PH* region is virtual (it is located in the region *FH*) and therefore the trajectories of the region *PH* cannot converge to it.

Figure 4 displays a phase plane of model (2) for *C*=0.15.

**Figure 4.**
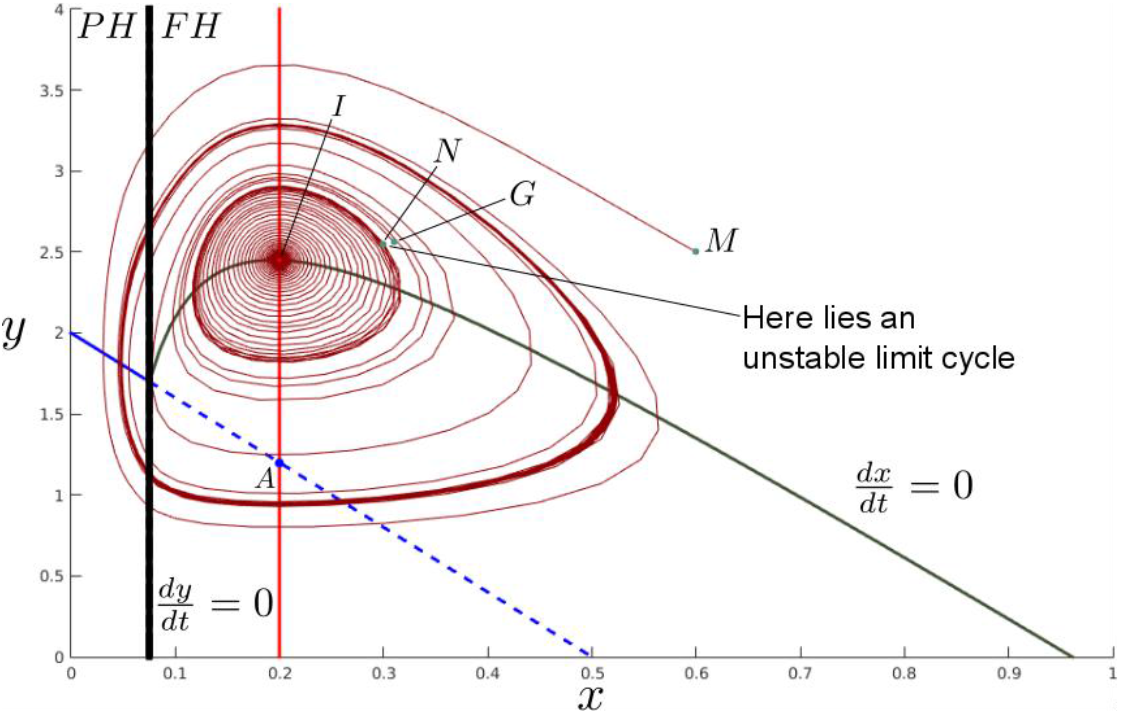
One phase plane of model (2) for *C* = 0.15. A higher value of *C* maintains the previous structure of equilibrium points of figure 3, but the combination of the vector fields of both structures generates an unstable limit cycle. 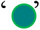 – initial conditions. Parameter values are the same as those of figure 3.

Figure 4 shows that a higher value of *C* maintains the previous structure of equilibrium points of figure 3, since point *A* of the *PH* region is still virtual and the equilibrium point *I* of the region *FH* is stable and real. Initial conditions at *M* lead to a stable limit cycle by means of convergent oscillations, while initial conditions at *G* lead to the same stable limit cycle by means of divergent oscillations. On the other hand initial conditions at *N* (very close to point *G*) lead to the real equilibrium point *I* of the *FH* region by means of an oscillatory convergence. This means that an unstable limit cycle passes between points *G* and *I*. Hence the combination of the vector fields of regions *PH* and *FH* can generate an unstable limit cycle encircling a locally stable equilibrium point and encircled by a stable limit cycle.

Figure 5 displays a phase plane of model (2) for *C*=0.35.

**Figure 5.**
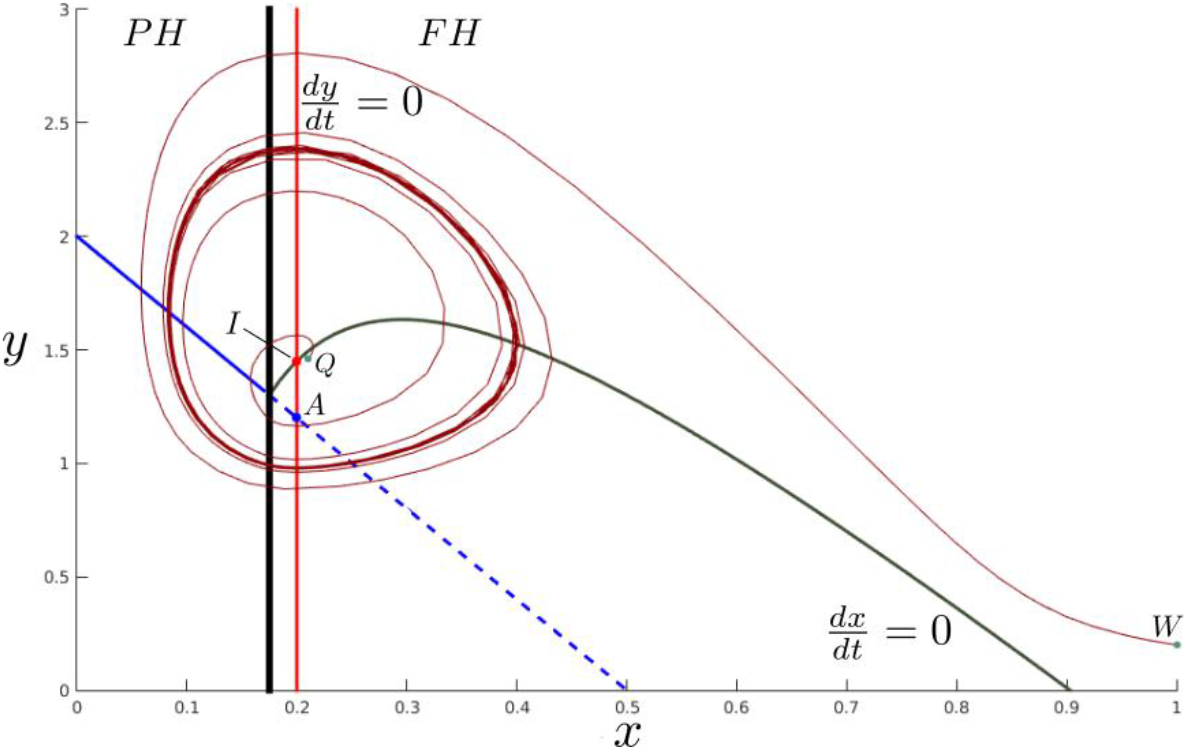
One phase plane of model (2) for *C* = 0.35. A higher value of *C* maintains the previous structures of equilibrium points of figures 3 and 4, but the combination of the vector fields of both structures generates a stable limit cycle. 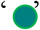 – initial conditions. Parameter values are the same as those of figure 3.

Figure 5 shows that a still higher value of *C* still maintains the previous two structures of equilibrium points of figures 3 and 4, but the combination of the vector fields now generates a stable limit cycle. Initial conditions at *Q* lead to the stable limit cycle by means of divergent oscillations, while initial conditions at *W* lead to the same stable limit cycle by means of convergent oscillations. Note also that the equilibrium point *I* of the *FH* region is still real but unstable now.

Figure 6 displays a phase plane of model (2) for *C*=0.405.

**Figure 6.**
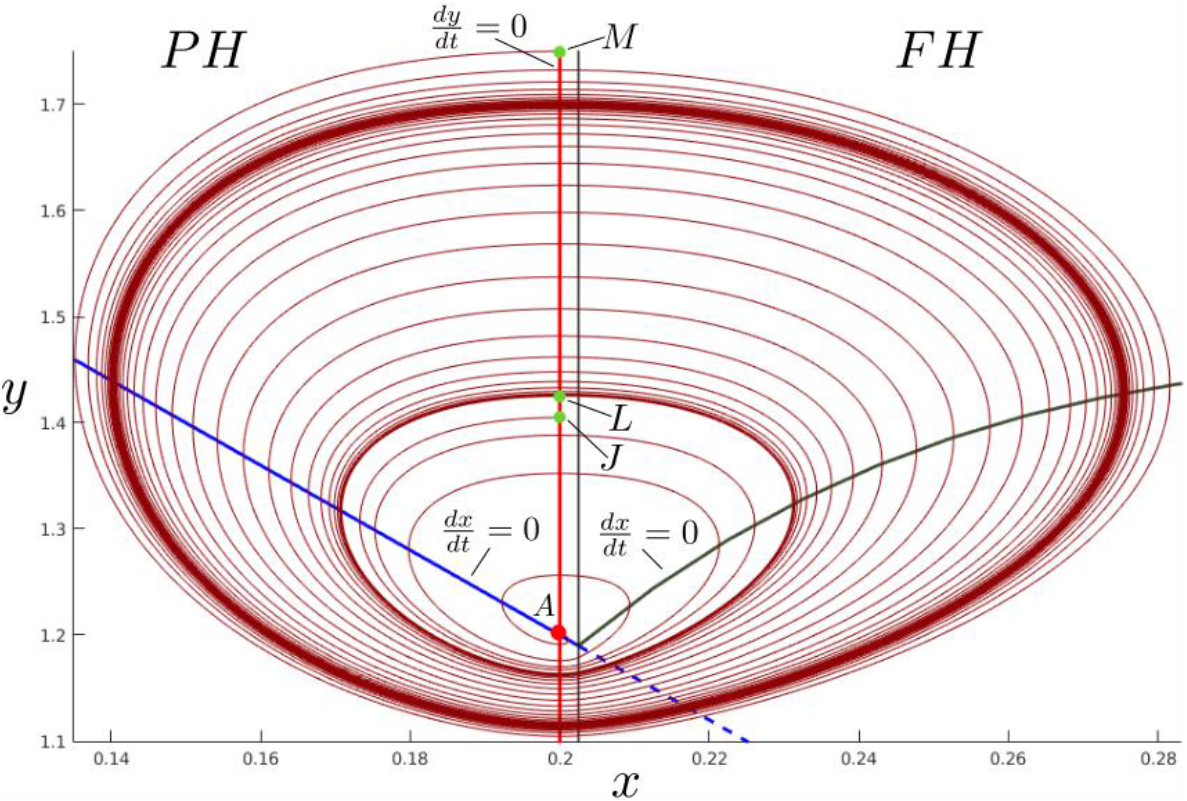
One phase plane of model (2) for *C* = 0.405. Increasing *C* still further, the equilibrium structure changes since the equilibrium point *A* of region *PH* becomes the only real and stable point. For initial conditions at *J* the trajectory converges to it, while at *L* and at *M* the trajectories converge to a stable limit cycle. 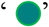 – initial conditions. Parameter values are the same as those of figure 3.

Figure 6 shows that increasing *C* still further, the equilibrium structure changes since the equilibrium point *A* of region *PH* becomes the only real and stable point. For initial conditions at *J* the trajectory converges to it, while at *L* and at *M* the trajectories converge to a stable limit cycle. This implies the existence of an unstable limit cycle between initial conditions *J* and *L* in the same way as in figure 4.

Figure 7 displays a phase plane of model (2) for *C*=0.5.

**Figure 7.**
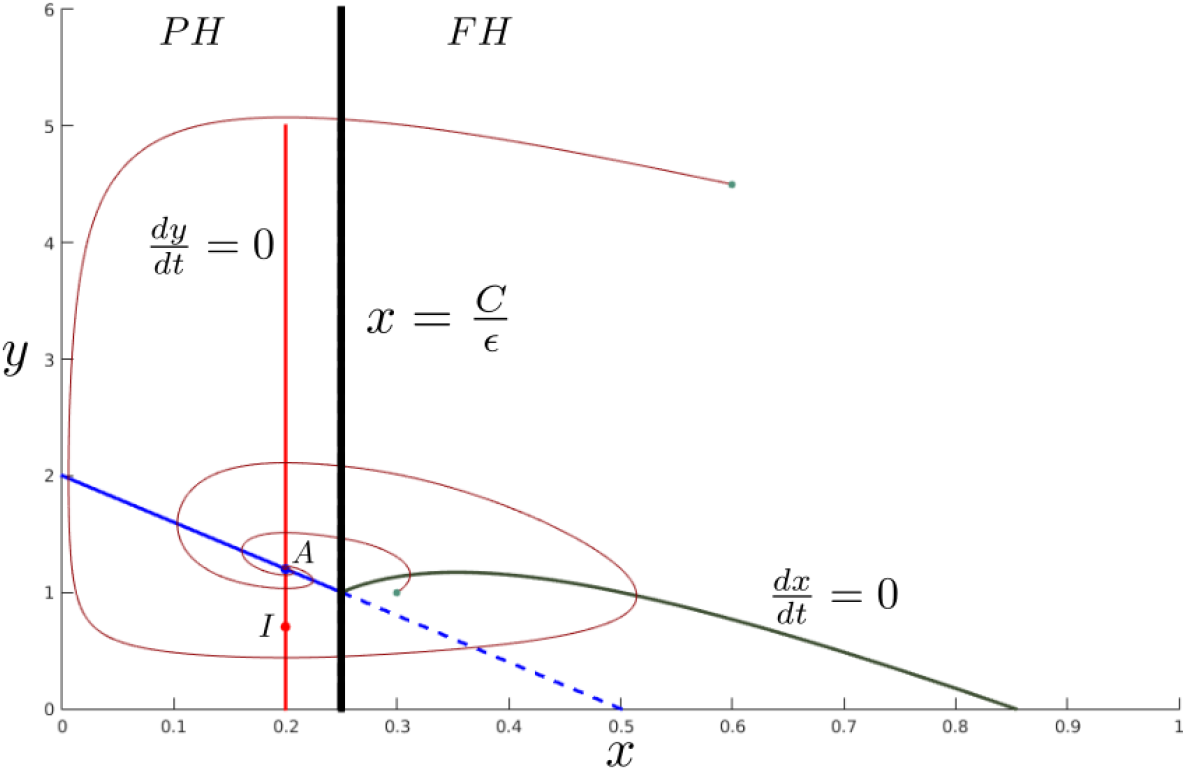
One phase plane of model (2) for *C* = 0.5. As in figure 6, increasing *C* further, the equilibrium structure remains the same since the equilibrium point *A* of region *PH* is still the only real and stable point, and in this case the trajectories converge to it. 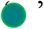 – initial conditions. Parameter values are the same as those of figure 3.

Figure 7 shows that increasing *C* further, the equilibrium structure remains the same since the equilibrium point *A* of region *PH* is still the only real and stable point, and in this case the simulated trajectory converges to it. In fact, from figure 6 we see that when 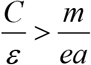, that is, when the switching line 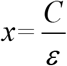 lies to the right of the intersection between the predator (*y*) isocline of the *PH* region 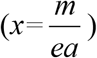 and the *x*-axis, the equilibrium point *A* of region *PH* will be the only real and stable point.

## 3 Discussion

With the purpose of understanding some aspects of the interplay between the behavior of exploited populations and the role of harvesting, this work dealt with a specific continuous time dynamical predator-prey model under prey harvest saturation.

As mentioned in the section Introduction, fishery has triggered many severe and unexpected consequences giving rise to communities that are more prone to cycles of booms and busts, and thus less manageable communities from the harvest standpoint. Our results pertaining to the phase planes of figures 3–7 come to reinforce this fact: prey harvesting saturation coupled with trophic interactions can produce complex dynamics rendering species management difficult. That is, the conjoint action of species interactions (in our case, predator-prey) and harvesting (in our case, prey harvest saturation) can push the population system past their tipping points. For instance, from figure 5 to 6, a slight increase of *C* changes a limit cycle into a limit cycle with a locally stable point in its interior. This dynamical change due to a slight disturbance certainly renders the population management more difficult since it depends now on population initial densities, requiring thus a fairly precise stock assessment.

At this stage, some comments are in order. The investigation of the separate influence of proportional harvest and constant harvest rate structures on some predator prey dynamics has already been performed by Brauer and Soudack (1979, 1982), showing a myriad of behaviors depending on the parameter values of *C, ε* and on population initial conditions. Hence, the coupling of these already investigated dynamics under the harvest saturation effect (expression (1)) together with the augmentation of the parameter *C* (within boundaries that guarantee a positive *x* isocline in the *FH* region) might probably generate complex dynamics as the ones previously shown here with their consequences to population management.

As mentioned before, in fisheries management models the term *min* (*εx C*) can represent a harvest saturation due, for instance, to engine limitations (Clark, 1976). Nonetheless, analyses of population mathematical models with fisheries described by non linear saturating functional responses (such as functional response type 2 and 3) are also encouraged (Mangel, 2006). In a similar context to the present study, the work of Costa et al. (2016) can be cast as a predator-prey system where both prey and predator are simultaneously harvested by a functional response type 3 (this setup could well describe the operation of international fishery fleets). It is shown that a slight increase in the size of a fishery fleet may bring about cycles of booms and busts, consequently rendering fishery less manageable in these cases. This result together with the ones presented here reinforces the importance of the interplay between species interactions and harvest strategies (Travis et al., 2014).

Given our focus on a specific harvest saturation function, it is important to cite some works in the population modeling literature that deals with some complex dynamics generated by the same saturating functional response type 1 used in this study. By means of computational simulations, Dubois and Closset (1975) demonstrated numerically the existence of one unstable and one stable limit cycle in the following predator-prey model

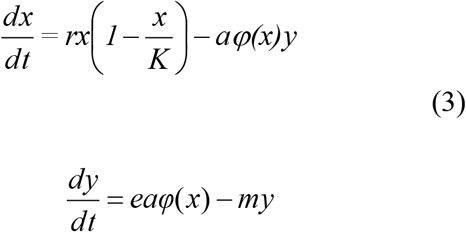

where *φ(x)*= min*(εx, C)* (they assume *ε*=1; the parameters in model (3) and (4) below have the same meanings as in model (2); see Seo and Kot (2008) for a mathematical demonstration of the existence of limit cycles in model (3)).

Adding fixed harvest rates *μ, γ* > 0 to the prey-predator model (3), that is,

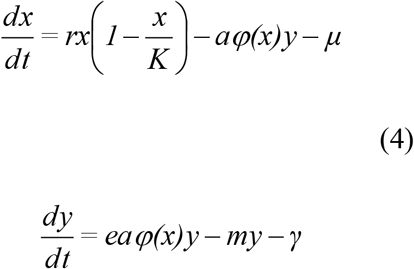

with *φ(x)* defined as in (3), Dai and Tang (1998) also found multiple limit cycles. These dynamical results together with the ones presented in this work show to some extent the built–in tendency to create multiple limit cycles when a saturated type 1 functional response is present in some predator–prey models. It is important to note, however, that models (3) and (4) possess a saturated type 1 functional response between a time variable predator (*y*) and the prey (*x*), while our model (2) possesses a saturated type 1 functional response between an additional predator with constant density (over time) and the prey (*x*).

To sum it up, encompassed by the importance of the interplay between species trophic interactions and harvest strategies in population management, the results of this study may serve as a contribution to the understanding of the influence that harvest functions with saturation may exert on the dynamics of exploited populations.

